## Supporting information including Figures S1-9 and Table S1 for "Quantification of *Salmonella enterica* serovar Typhimurium Population Dynamics in Murine Infection Using a Highly Diverse Barcoded Library"

**This file includes:**

Figures S1 to S9
Table S1
SI References

**Other supporting materials for this manuscript include the following:**

Dataset S1

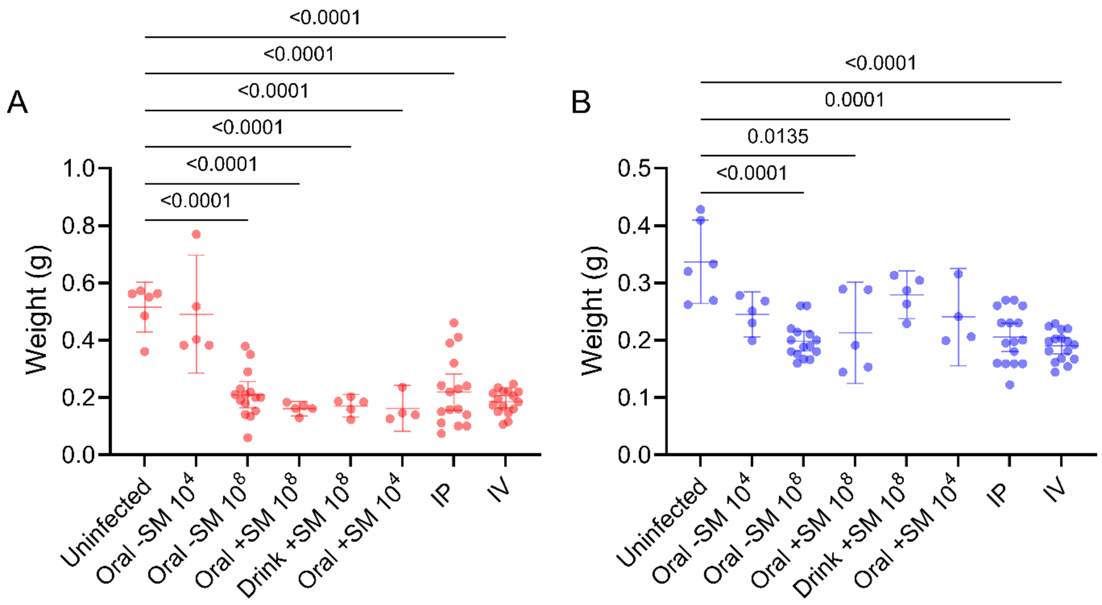


**Fig. S1. Weight of cecum (A) and colon (B) after different infection schemes.** Mann-Whitney tests were used for statistical analyses.

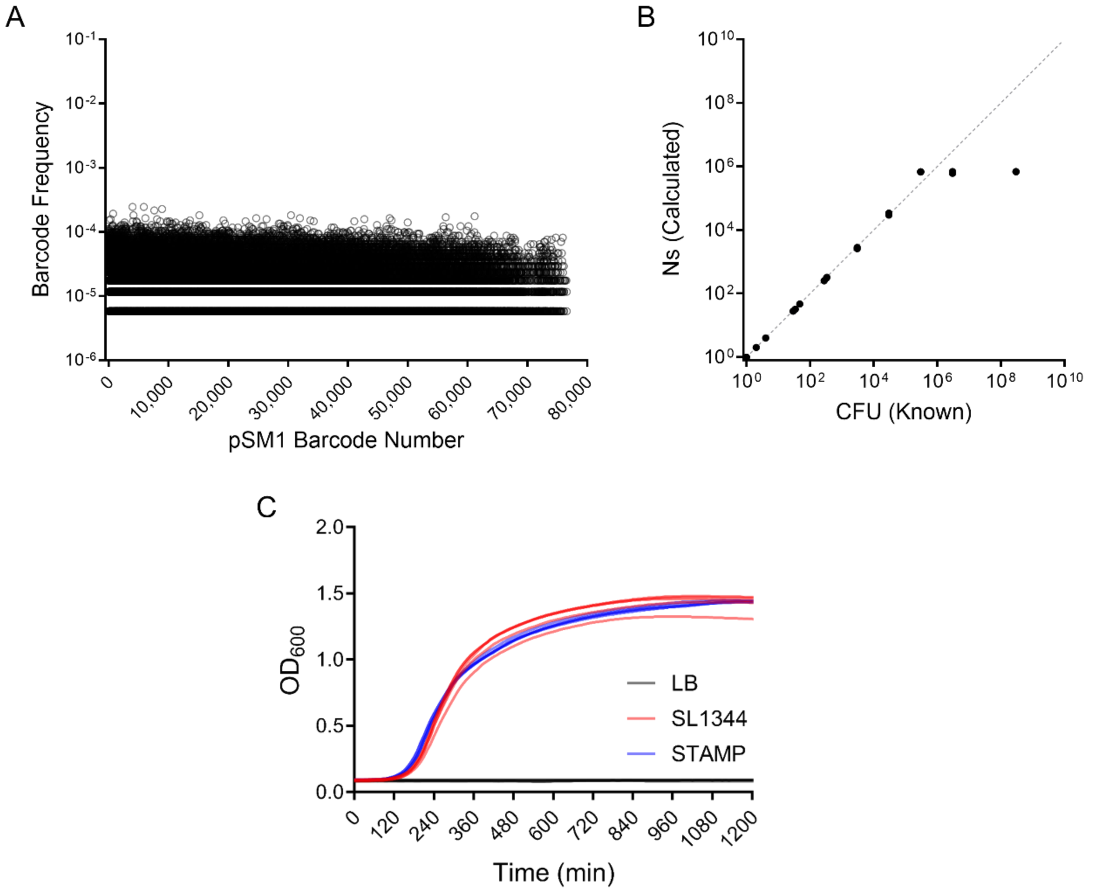


**Fig. S2. *S*. Typhimurium barcoded library diversity and standard curve. A)** Barcodes in *S* Typhimurium library were mapped to the barcodes in the donor library (pSM1) and the frequency of barcodes in the *S*. Typhimurium library were distributed relatively evenly. **B)** Calibration curve indicating the library has an Ns resolution limit of ~7x$10^5$. **C)** Growth of barcoded STAMP library in LB supplemented with SM is not different than the parent strain (SL1344). Significance was tested with Mann-Whitney test and growth curves were performed 5 times. Curves displayed with 50% opacity.

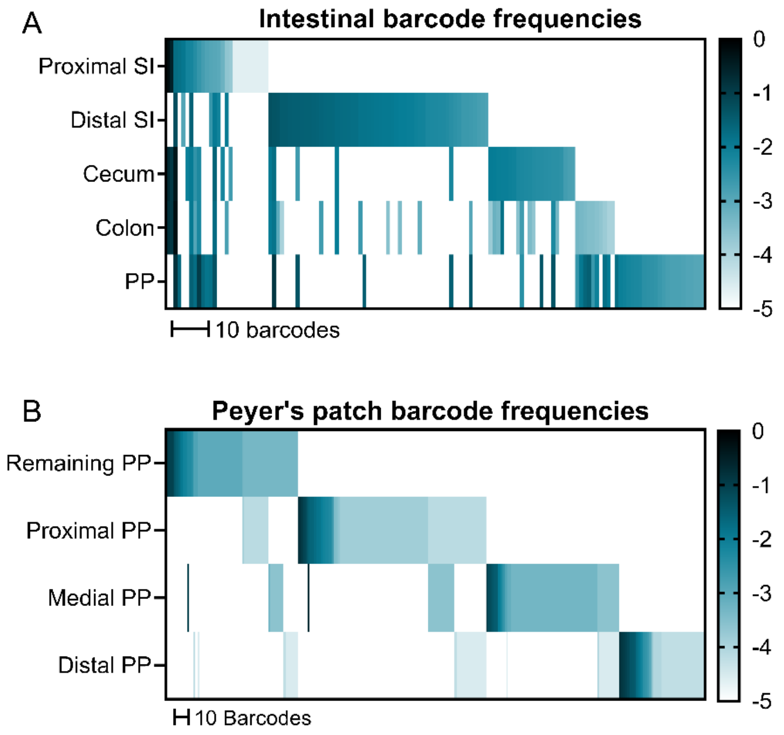


**Fig. S3. Barcodes from different regions of the intestine and Peyer's patches from a single animal are largely distinct. A)** Barcode frequencies of different regions of the intestine and **B)** individual Peyer's patches (PP) of an untreated mouse after orogastric gavage of *S*. Typhimurium. "Remaining PP" sample contains the PP from the animal not taken for individual analysis. Frequencies of barcodes are displayed as heatmaps (each box represents a barcode and darker colors represent increased frequency within the sample) and sorted by abundance in each sample sequentially.

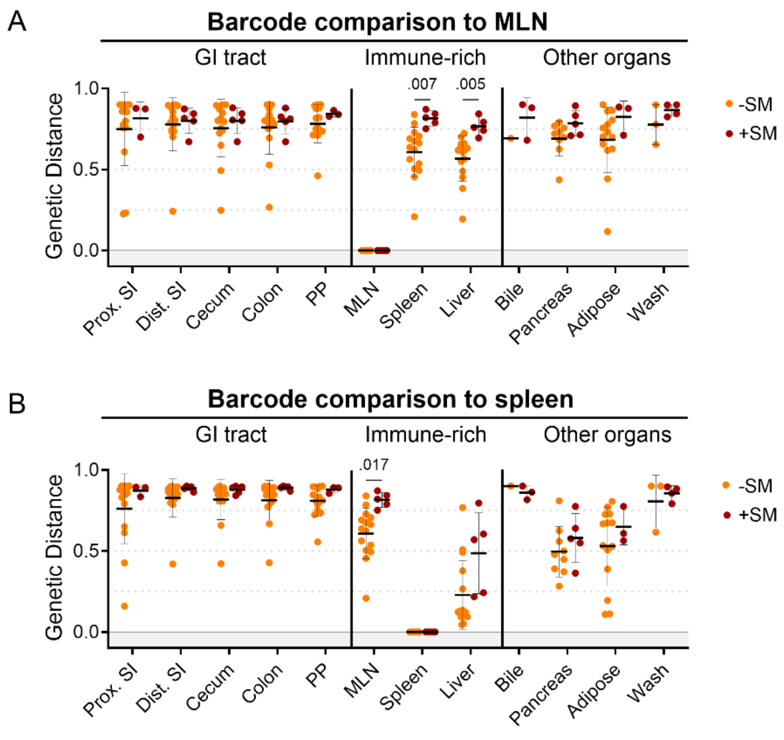


**Fig. S4. *S*. Typhimurium disseminates to the MLN and spleen prior to substantial replication in the intestine**. Genetic distance from the MLN (**A**) and spleen (**B**) after orogastric inoculation of *S*. Typhimurium with (+SM) or without (-SM) streptomycin pretreatment. -SM n =16 (8 males & 8 females), +SM n = 5 (female). Mann-Whitney tests were used for statistical analyses. Values with p < 0.1 shown. Abbreviations: GI, gastrointestinal; MLN, mesenteric lymph node; PP, Peyer's patches; SI, small intestine.

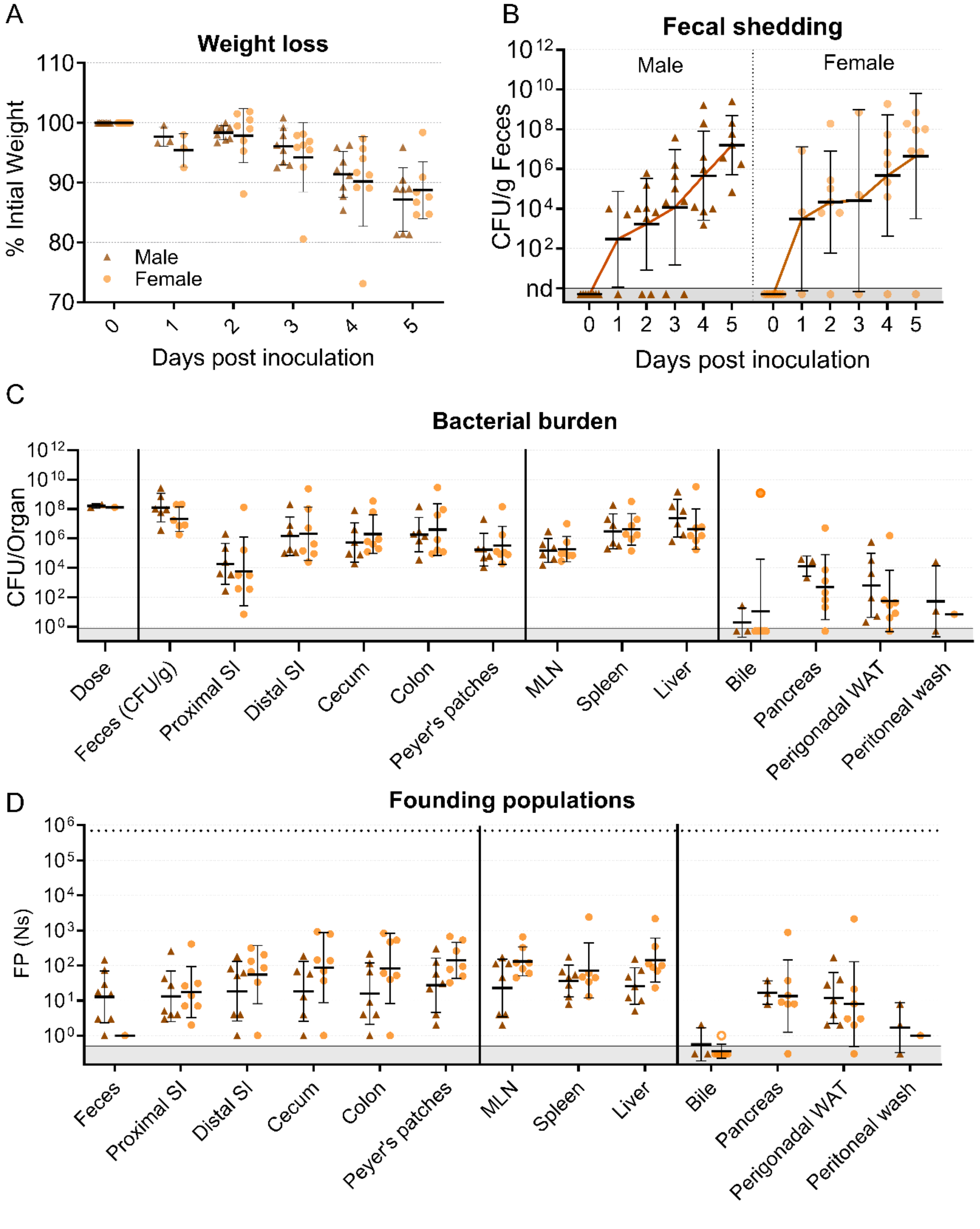


**Fig. S5. Sex-disaggregated data in mice inoculated through orogastric gavage. A)** Percentage of initial weight over time; means and standard deviations shown. **B)** Bacterial burden in feces. **C)** Bacterial burden in organs and fluids, open circles indicate when the whole gallbladder was taken instead of bile. **D)** Founding populations (Ns) in organs and fluids. Geometric means and geometric standard deviations are used unless otherwise noted. -SM female n = 8, -SM male n = 8

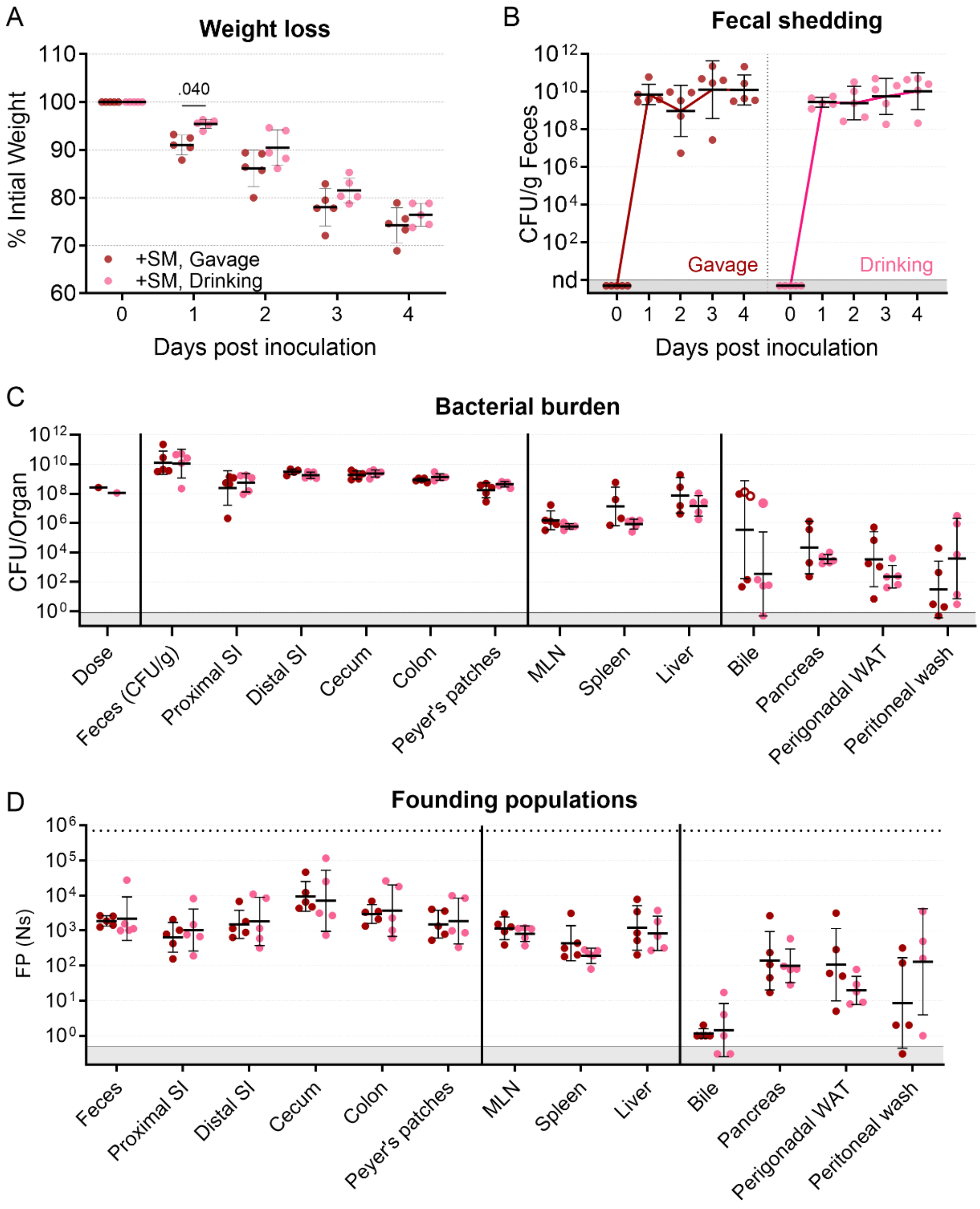


**Fig. S6. Extended data for streptomycin pretreated mice inoculated through orogastric gavage and drinking. A)** Percentage of initial weight over time; means and standard deviations shown. **B)** Bacterial burden in feces. **C)** Bacterial burden in organs and fluids, open circles indicate when the whole gallbladder was taken instead of bile. **D)** Founding populations (Ns) in organs and fluids. Geometric means and geometric standard deviations are used unless otherwise noted. +SM, Gavage female n = 5, and +SM, Drinking female n = 5. Mann-Whitney tests were used for statistical analyses. Values with $p < 0.1$ shown.

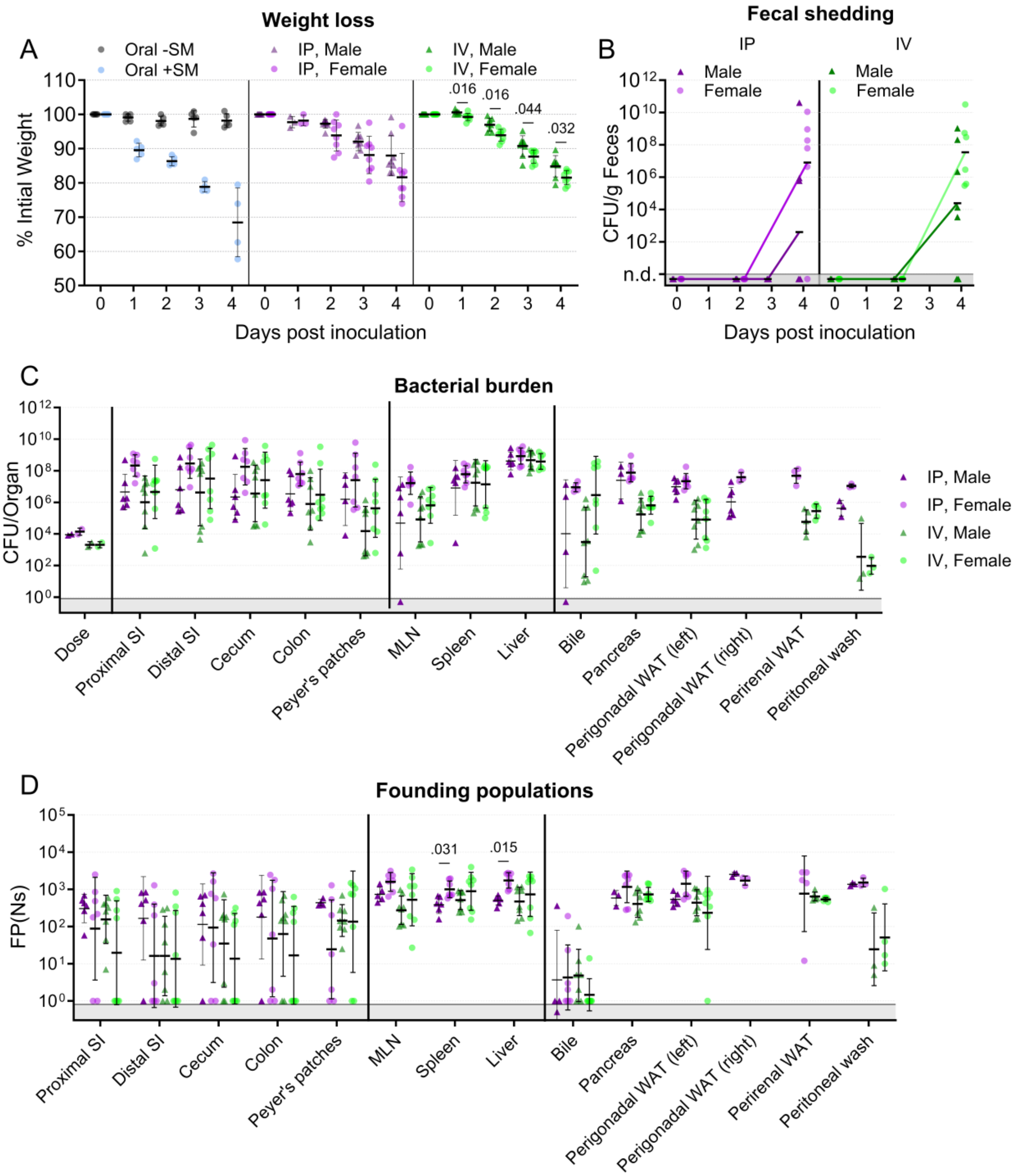


**Fig. S7. Sex-disaggregated data in mice after inoculation with $10^3$-$10^4$ CFU *S.* Typhimurium through different routes. A)** Percentage of initial weight over time; means and standard deviations shown. **B)** Sex-disaggregated bacterial burden in feces. **C)** Bacterial burden in organs and fluids. **D)** Founding populations (Ns) in organs and fluids. Geometric means and geometric standard deviations are used unless otherwise noted. Oral -SM female n = 5, Oral +SM n = 5, IP male n = 8, IP female n = 8, IV male n = 8, and IV female n = 8. Statistical analysis is a Mann-Whitney test. Values with $p < 0.1$ are shown.

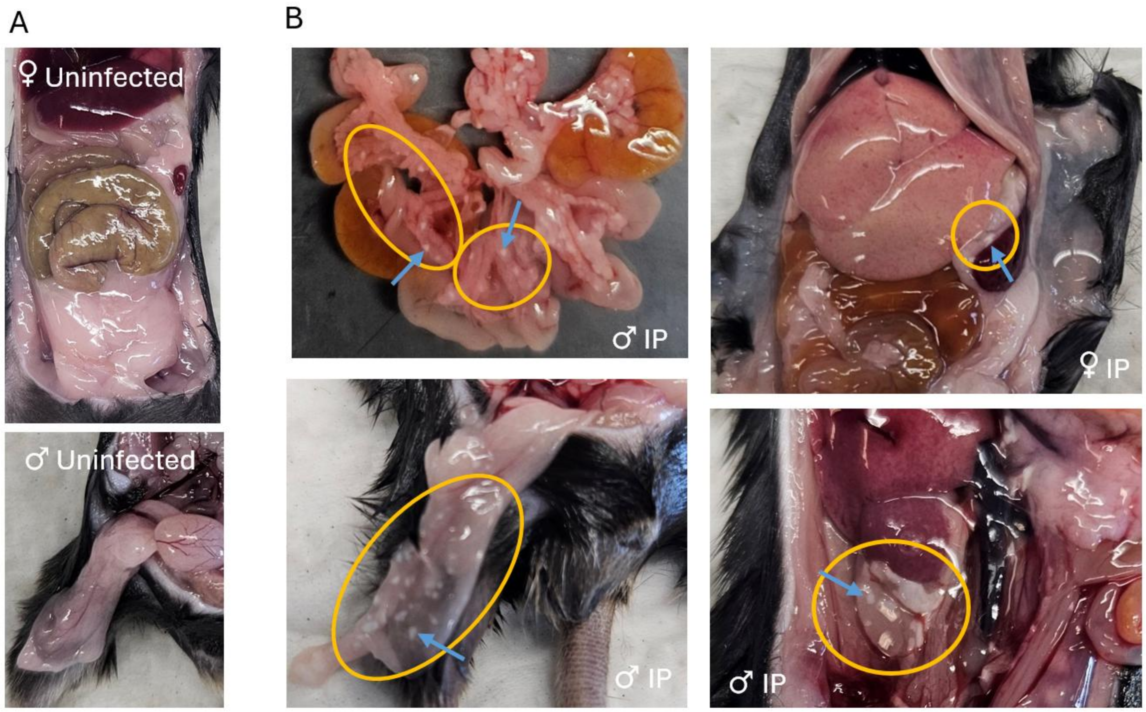


**Fig. S8. Pathology observed in mouse adipose tissue. A)** Images of uninfected adipose. **B)** Images of white spots on mesenteric (top left), upper mesenteric (top right), perigonadal (bottom left) and perirenal (bottom right) adipose in mice. Yellow circles outline areas with pathology. Blue arrows point to focal lesions. Abbreviations: IV, intravenous; IP, intraperitoneal

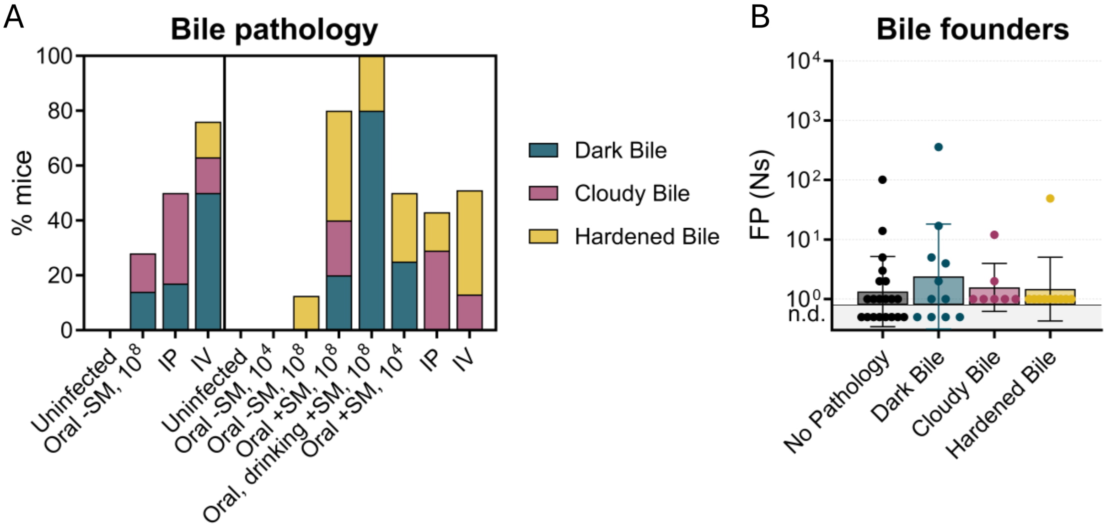


**Fig. S9. Bile pathology correlates with the bile re-seeding pathway. A)** Percentage of observed biliary pathology. **B)** Founding population of bile disaggregated by biliary pathology. Abbreviations: IV, intravenous; IP, intraperitoneal; SM, streptomycin; SI, small intestine

**Table S1.** Primer sequences for PCR.

| Name | Sequence |
|---|---|
| Forward Primers | |
| var21 | AATGATACGGCGACCACCGAGATCTACACTCTTTCCCTACACGACGCTCTTCCGATCTAATGATGGG TTAAAAAGGATCGATCC |
| var22 | AATGATACGGCGACCACCGAGATCTACACTCTTTCCCTACACGACGCTCTTCCGATCTATGCGATGG GTTAAAAAGGATCGATCC |
| var23 | AATGATACGGCGACCACCGAGATCTACACTCTTTCCCTACACGACGCTCTTCCGATCTTGCACGATG GGTTAAAAAGGATCGATCC |
| var24 | AATGATACGGCGACCACCGAGATCTACACTCTTTCCCTACACGACGCTCTTCCGATCTTCATTCGAT GGGTTAAAAAGGATCGATCC |
| var25 | AATGATACGGCGACCACCGAGATCTACACTCTTTCCCTACACGACGCTCTTCCGATCTGAATCGAGA TGGGTTAAAAAGGATCGATCC |
| var26 | AATGATACGGCGACCACCGAGATCTACACTCTTTCCCTACACGACGCTCTTCCGATCTGTCAACTTG ATGGGTTAAAAAGGATCGATCC |
| var27 | AATGATACGGCGACCACCGAGATCTACACTCTTTCCCTACACGACGCTCTTCCGATCTCGGCGTGGC GATGGGTTAAAAAGGATCGATCC |
| var28 | AATGATACGGCGACCACCGAGATCTACACTCTTTCCCTACACGACGCTCTTCCGATCTCCTGTACCT TGATGGGTTAAAAAGGATCGATCC |
| Reverse Primers | |
| AD001 | CAAGCAGAAGACGGCATACGAGATCGTGATGTGACTGGAGTTCAGACGTGTGCTCTTCCGATCAGAT CCTTGGCGGCAAGAAA |
| AD002 | CAAGCAGAAGACGGCATACGAGATACATCGGTGACTGGAGTTCAGACGTGTGCTCTTCCGATCAGAT CCTTGGCGGCAAGAAA |
| AD003 | CAAGCAGAAGACGGCATACGAGATGCCTAAGTGACTGGAGTTCAGACGTGTGCTCTTCCGATCAGAT CCTTGGCGGCAAGAAA |
| AD004 | CAAGCAGAAGACGGCATACGAGATTGGTCAGTGACTGGAGTTCAGACGTGTGCTCTTCCGATCAGAT CCTTGGCGGCAAGAAA |
| AD005 | CAAGCAGAAGACGGCATACGAGATCACTGTGTGACTGGAGTTCAGACGTGTGCTCTTCCGATCAGAT CCTTGGCGGCAAGAAA |
| AD006 | CAAGCAGAAGACGGCATACGAGATATTGGCGTGACTGGAGTTCAGACGTGTGCTCTTCCGATCAGAT CCTTGGCGGCAAGAAA |
| AD007 | CAAGCAGAAGACGGCATACGAGATGATCTGGTGACTGGAGTTCAGACGTGTGCTCTTCCGATCAGAT CCTTGGCGGCAAGAAA |
| AD008 | CAAGCAGAAGACGGCATACGAGATTCAAGTGTGACTGGAGTTCAGACGTGTGCTCTTCCGATCAGAT CCTTGGCGGCAAGAAA |
| AD009 | CAAGCAGAAGACGGCATACGAGATCTGATCGTGACTGGAGTTCAGACGTGTGCTCTTCCGATCAGAT CCTTGGCGGCAAGAAA |
| AD010 | CAAGCAGAAGACGGCATACGAGATAAGCTAGTGACTGGAGTTCAGACGTGTGCTCTTCCGATCAGAT CCTTGGCGGCAAGAAA |
| AD011 | CAAGCAGAAGACGGCATACGAGATGTAGCCGTGACTGGAGTTCAGACGTGTGCTCTTCCGATCAGAT CCTTGGCGGCAAGAAA |
| AD012 | CAAGCAGAAGACGGCATACGAGATTACAAGGTGACTGGAGTTCAGACGTGTGCTCTTCCGATCAGAT CCTTGGCGGCAAGAAA |
| AD013 | CAAGCAGAAGACGGCATACGAGATTTGACTGTGACTGGAGTTCAGACGTGTGCTCTTCCGATCAGAT CCTTGGCGGCAAGAAA |
| AD014 | CAAGCAGAAGACGGCATACGAGATGGAACTGTGACTGGAGTTCAGACGTGTGCTCTTCCGATCAGAT CCTTGGCGGCAAGAAA |
| AD015 | CAAGCAGAAGACGGCATACGAGATTGACATGTGACTGGAGTTCAGACGTGTGCTCTTCCGATCAGAT CCTTGGCGGCAAGAAA |
| AD016 | CAAGCAGAAGACGGCATACGAGATGGACGGGTGACTGGAGTTCAGACGTGTGCTCTTCCGATCAGAT CCTTGGCGGCAAGAAA |
| AD018 | CAAGCAGAAGACGGCATACGAGATGCGGACGTGACTGGAGTTCAGACGTGTGCTCTTCCGATCAGAT CCTTGGCGGCAAGAAA |

| | |
|---|---|
| AD019 | CAAGCAGAAGACGGCATACGAGATTTTCACGTGACTGGAGTTCAGACGTGTGCTCTTCCGATCAGATCCTTGGCGGCAAGAAA |
| AD020 | CAAGCAGAAGACGGCATACGAGATGGCCACGTGACTGGAGTTCAGACGTGTGCTCTTCCGATCAGATCCTTGGCGGCAAGAAA |
| AD021 | CAAGCAGAAGACGGCATACGAGATCGAAACGTGACTGGAGTTCAGACGTGTGCTCTTCCGATCAGATCCTTGGCGGCAAGAAA |
| AD022 | CAAGCAGAAGACGGCATACGAGATCGTACGGTGACTGGAGTTCAGACGTGTGCTCTTCCGATCAGATCCTTGGCGGCAAGAAA |
| AD023 | CAAGCAGAAGACGGCATACGAGATCCACTCGTGACTGGAGTTCAGACGTGTGCTCTTCCGATCAGATCCTTGGCGGCAAGAAA |
| AD025 | CAAGCAGAAGACGGCATACGAGATATCAGTGTGACTGGAGTTCAGACGTGTGCTCTTCCGATCAGATCCTTGGCGGCAAGAAA |
| AD027 | CAAGCAGAAGACGGCATACGAGATAGGAATGTGACTGGAGTTCAGACGTGTGCTCTTCCGATCAGATCCTTGGCGGCAAGAAA |

**Dataset S1 (separate file).** Barcode counts for all samples in manuscript.
